# Multi-omics Integrative Analysis of Acute and Relapsing Malaria in a Non-Human Primate Model of *P. vivax* infection

**DOI:** 10.1101/564195

## Abstract

Systems-scale analysis of multiple layers of molecular and cellular data has significant potential for providing novel insights into malaria pathology and immunity. We present here a unique longitudinal multi-omics dataset encompassing *Macaca mulatta* blood and bone marrow responses to infection by *Plasmodium cynomolgi*, a non-human primate (NHP) parasite species used to model *P. vivax* malaria acute and relapsing infections in humans. We analyzed relationships across multiple biological layers using a mutual information-based machine learning approach to integrate heterogeneous longitudinal datasets and constructed an atlas of multi-omics relatedness networks (MORNs). Using this technique, we were able to detect signatures that defined both acute and relapsing infections. Importantly, relapse infections could be distinguished from both acutely-infected and uninfected NHP, suggesting that the host-parasite interactions during relapses are unique compared to acute *Plasmodium* infections. To our knowledge, this is the first report of large-scale, longitudinal multi-omics analysis of malaria in any system. This dataset, along with the method used to analyze it, provides a unique resource for the malaria research community and demonstrates the power of longitudinal infection study designs, NHP model systems and integrative multi-omics analyses.

## Introduction

*Plasmodium vivax* infections cause substantial morbidity, with an estimated 8.5 million infections each year and 2.85 billion people at risk of disease (Guerra et al., 2010). One of the unique aspects of *P. vivax* biology is its ability to form dormant liver-stage forms. These are known as hypnozoites and can persist and exit dormancy causing relapsing blood-stage infections weeks, months or years after an initial blood-stage infection has been cleared. Relapses are thought to be responsible for most blood-stage infections in *P. vivax* endemic areas (Adekunle et al., 2015), although there is difficulty in determining this. Regardless, it is widely accepted that relapsing *P. vivax* infection possesses the potential to continue transmission and cause repeated bouts of malaria. A better understanding of relapse infection biology is thus critical for current malaria control and eradication efforts. The *Plasmodium cynomolgi* – rhesus macaque (*Macaca mulatta*) infection model for vivax malaria holds great potential for studying host responses and pathogenesis during acute and relapsing Plasmodium infections (Joyner et al., 2016). Macaque infections allow for more tractable longitudinal sampling of peripheral blood (PB) and bone marrow (BM) than in human studies, and macaque and human immune systems are highly similar, making this an excellent model system.

Here, we present a unique longitudinal dataset from a multi-omics study designed to investigate host responses in *M. mulatta* and compare *P. cynomolgi* acute infections with relapse infections. We integrated transcriptomics, metabolomics, and lipidomics data, and synthesized them with immunophenotyping using a unique and intuitive mutual information-based network analysis approach. This approach distinguished between acute infections, relapse infections, and uninfected NHP in an unsupervised manner to an extent that was not possible based on the individual data types alone. Furthermore, relapse infections could be distinguished from uninfected NHP. Functional enrichment for a number of the identified molecular networks was validated using existing transcriptional datasets from human *Plasmodium* infections, supporting the generalizability of our results.

## Results

To gain a systems-scale understanding of possible host responses in vivax malaria, we performed transcriptomics, untargeted metabolomics, and targeted lipidomics analyses using bone marrow (BM) and peripheral blood (PB) samples acquired longitudinally across a 100-day infection of four *M. mulatta* infected with *P. cynomolgi* (Figure 1A). We integrated these multi-omic datasets together and with other datasets generated from the same samples, including immunophenotyping, cytokine profiling, and parasite transcriptional profiling. Samples spanned pre-infection, acute primary infection, post-peak, and relapse infection time points. Comprehensive details on the longitudinal experimental design, clinical parameters relating to parasitemia and anemia, as well as the sampling time points have been described previously (Joyner et al., 2016).

**Figure 1.**
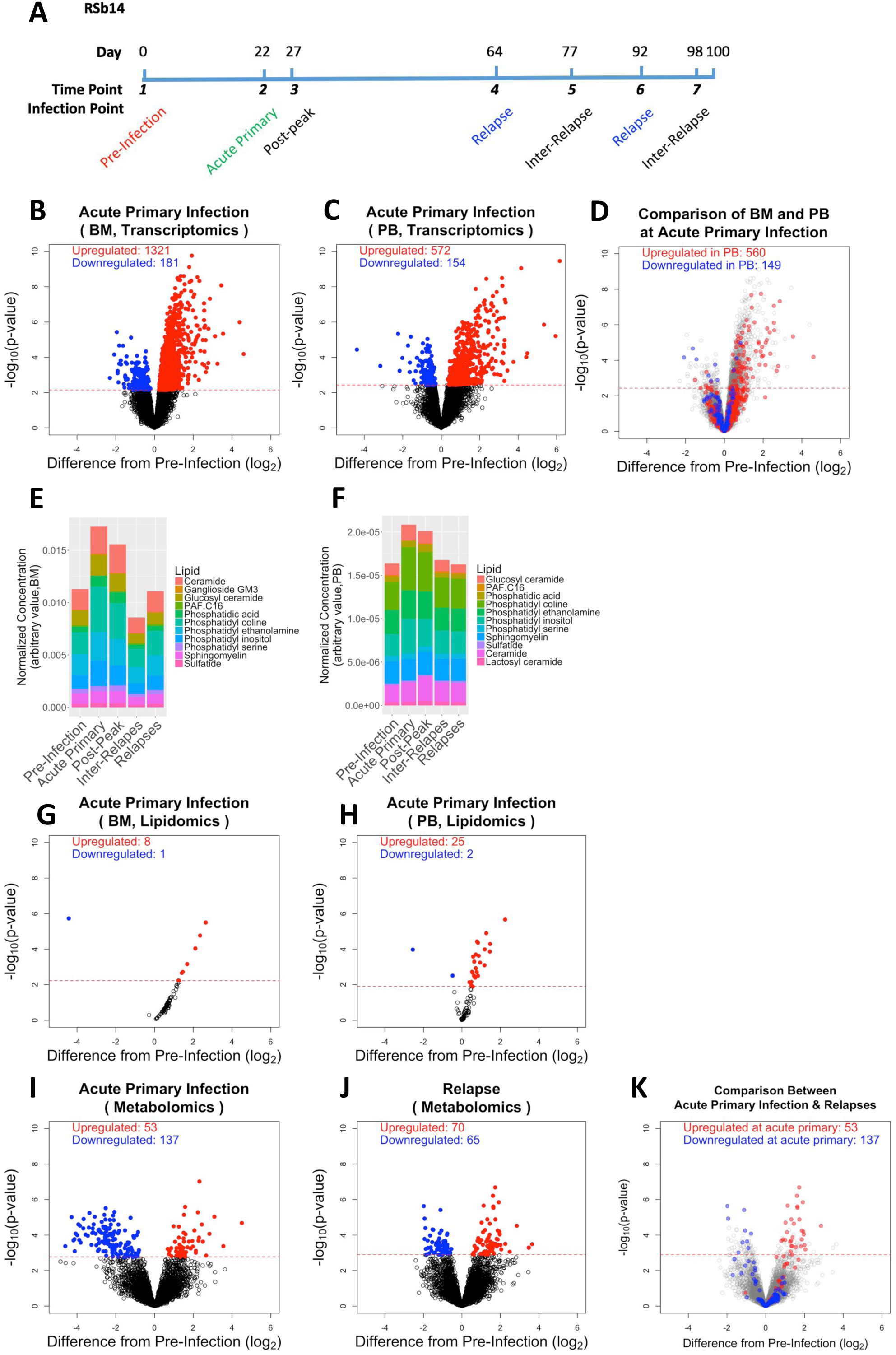
Altered omics features at acute primary infection. (A) Representative schematics for experiment design for one macaque (RSb14). Over 100 days, peripheral blood and bone marrow aspirate samples were collected at 7 time points, including pre-infection (day 0), acute primary infection (day 22), post-peak (day 27), relapses (day 64, 92) and inter-relapse (day 77, 98). (B) Differential expression (DE) analysis shows skewed upregulation of gene expression at acute primary infection in BM. Red dots represent upregulated genes; blue dots represent downregulated genes. x axis shows fold change; y axis shows p-value of significance in negative log10. Dotted red line indicates a significance threshold of 0.05 (FDR, Benjamini-Hochberg test). (C) DE analysis shows skewed upregulation of gene expression at acute primary infection in PB. (D) DEGs in BM, with colored dots overlaid based on DE status of the same genes in PB at acute primary infection. Red dots represent genes upregulated in PB, with their fold change in BM being plotted; blue dots represent genes downregulated in PB, with their fold change in BM being plotted. (E) Lipid subclasses measured in BM. Stacked colored bars represent the composition of lipid subclasses at each infection point. (F) Lipid subclasses measured in PB. (G-H) Differential accumulation (DA) analysis shows skewed upregulation of lipids at acute primary infection in BM (G) and in PB (H). (I) DA analysis shows skewed downregulation of metabolite abundance at acute primary infection. (J) DA analysis shows differential abundance of metabolic features at relapses. (K) Metabolic features with differential abundance at relapses, with colored dots overlaid representing differential abundance status of the same features at acute primary infection. Red dots represent features upregulated at acute primary infection, with their fold change in relapse being plotted; blue dots represent features downregulated at acute primary infection, with their fold change in relapse being plotted.

### Acute and relapse infections have differing profiles

Clustering features across all omics data and organizing samples by infection time point shows that acute infection was characterized by the broadest scope and greatest magnitude of changes in omic profiles (Figure S1A), a result that was expected given the lower parasite burden during relapse infection (Joyner et al., 2016). Acute infections led to significantly upregulated transcripts for many genes (differentially expressed genes; DEG) compared to baseline (Figure 1B-C). Very few DEGs were identified at relapse as compared to baseline, with none noted in the BM (Figure S1B-C). This was consistent with lipidomics profiles which showed strong changes only during acute infection and immediately after the peak of parasitemia, but not in relapse infections (Figure 1E-H). It also agreed with immune profiling where relapsing individuals looked similar to uninfected individuals (Figure 2, discussed in greater depth below). Untargeted high resolution metabolomics (HRM) analysis of plasma yielded over 5,000 metabolic features after filtering. Surprisingly, metabolomics data showed a very low subset of features with significant changes at any given time point (only around 4%). In contrast with the transcriptomics data, HRM data showed a similar number of overall features with differential abundance at acute infection and relapse time points compared to pre-infection (Figure 1I-J). Furthermore, the direction of changes in HRM data was more conserved between acute infection and relapse (Figure 1K). Thus, metabolomics profiling with univariate analysis can distinguish infected from uninfected individuals regardless of the status of the infection, whereas transcriptomic, lipidomic and immune profiling were only able to identify differences during an acute infection.

**Figure 2.**
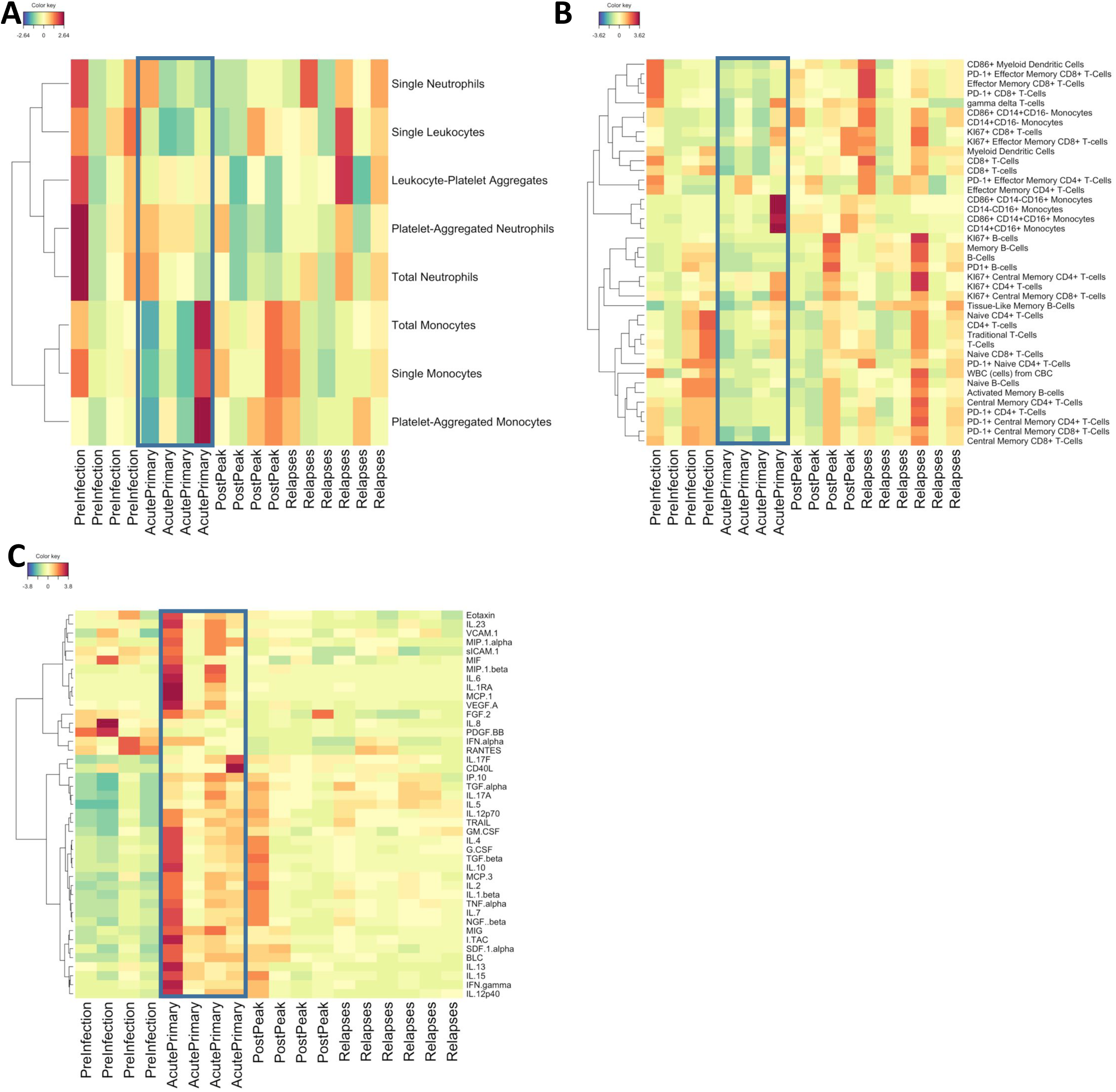
Immunological profiling (A) Profiling of innate immune cell populations, with cell types hierarchically clustered. Data are median-subtracted and log-scaled. (B) Profiling of adaptive immune cell populations. (C) Profiling of plasma cytokines.

To determine whether measured metabolomic profiles during relapse infection were distinct from those during acute infection or just lower intensity versions of the same changes, we identified the major axes of variation in individual data types using principal component analysis (PCA). For most data types, the first two principal components were insufficient to separate between time-points in an unsupervised fashion (Figure 3A), suggesting that the dominant modes of variation in the dataset reflect a non-temporal relationship between samples or biological noise. Strikingly, PCA using all variables measured across all data types for either BM or PB showed a significant improvement in the ability to distinguish between infection time points (Figure 3B), supporting the utility of multi-omics analysis. The improvement in sample classification when combining data types suggested that integration of the varied datasets might yield improved molecular signatures.

**Figure 3.**
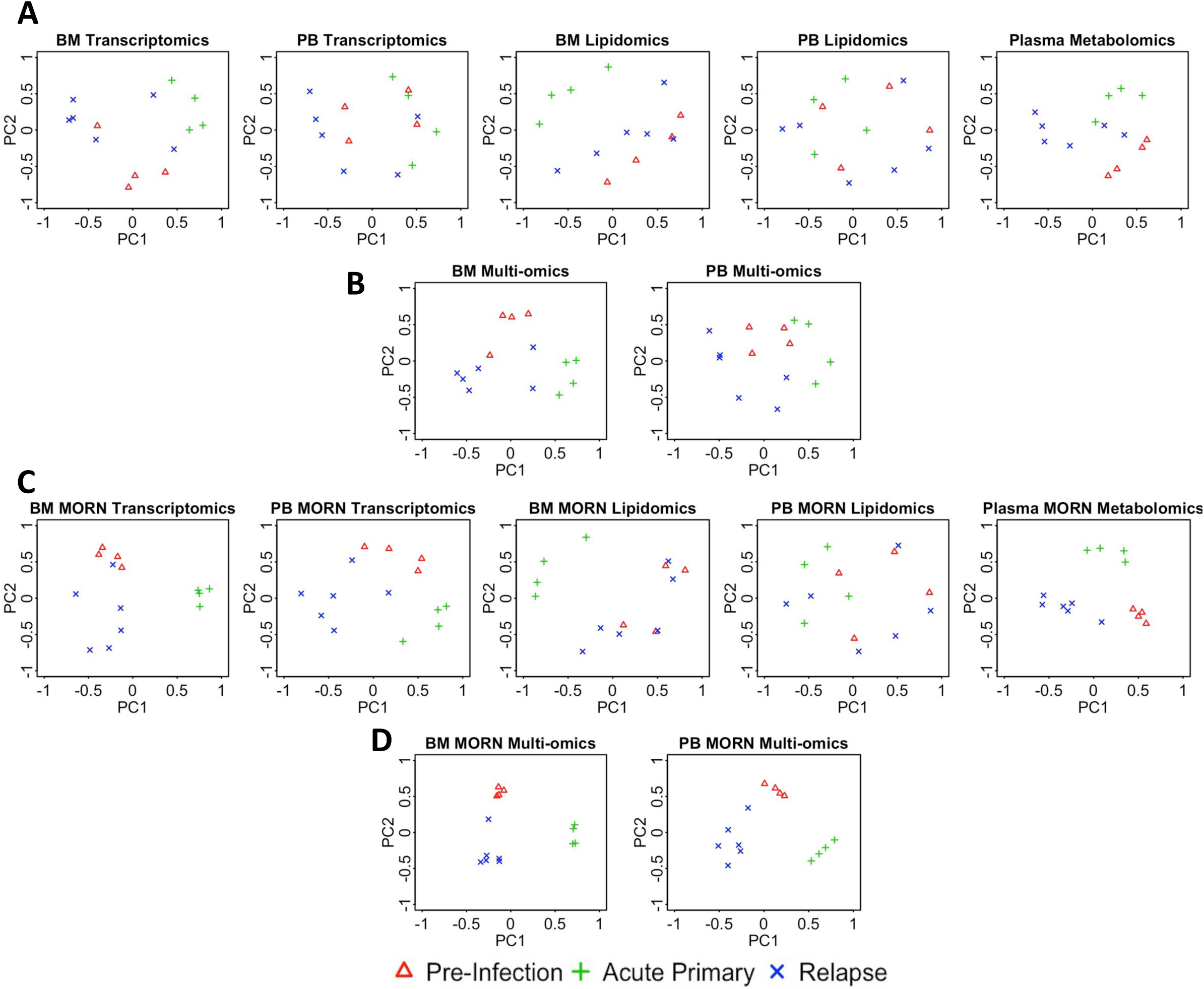
Unsupervised analysis of individual omics dataset and multi-omics dataset profiles. (A) PCA of individual omics datasets, showing minimal separation of samples based on infection points. First and second principal component projections on x and y axes, respectively, yield minimal clustering of samples at pre-infection (red), acute primary infection (green), and relapse (blue) infection points. (B) PCA of all omics datasets together in PB and in BM shows improved separation of infection points compared to analysis of individual omics datasets. (C) PCA of individual omics datasets only with features selected based on cross-omics mutual information shows markedly improved clustering of samples by infection points. (D) PCA of all omics features selected based on cross-omics mutual information in PB and in BM shows complete separation of sample clusters based on infection points.

### Bone marrow differs from peripheral cells

BM disturbances are known to be a major factor in malaria. Having determined that acute infection led to marked changes at all molecular levels measured, we determined whether there were significant differences in the PB compared to the BM. After filtering out low expression genes, we measured the transcription of 10,487 and 9,735 genes in the PB and BM, respectively. We observed a large overlap between the two compartments with a notable skew towards upregulation in both compartments. Although more DEGs were observed in BM (Figure 1B) compared with PB samples (Figure 1C) at acute infection compared to baseline, only a small number of shared DEGs were identified, implying activation of different transcriptional programs in the two compartments (Figure 1D). At the acute infection time point, Gene Ontology (GO) enrichment analysis of DEGs with ToppFun (Chen et al., 2009) identified distinct functional categories involved in PB and BM. Top enriched GO biological processes in PB were mostly related to the immune response and immune cell number in the periphery (**Table S1**), consistent with more pronounced immune responses at acute infection. In contrast, enrichment analysis of the BM indicated involvement of responses to unfolded protein, RNA processing, and regulation of the stress response (**Table S2**).

To make sense of the PB cellular dynamics driving the transcriptomic results across infection time-points, we performed longitudinal profiling of innate (Figure 2A) and adaptive (Figure 2B) immune cell subsets in PB alongside quantification of plasma cytokines (Figure 2C). The total number of leukocytes in the periphery was lower during acute infection and a large proportion of circulating cells displayed a memory phenotype (Figure 2A-B). This may have occurred due to trafficking to, and retention in, lymphoid organs and may be a significant feature of acute infection as repeatedly demonstrated in the lung and liver during mouse models of *Plasmodium* infection (Lagasse et al., 2016; Mimche et al., 2015). Indeed, this idea is consistent with cytokine levels in PB that generally increased at acute infection compared to baseline, demonstrating an elevated response (Figure 3C). At the post-peak and the relapse time points cell numbers returned to the levels observed prior to infection.

Targeted lipidomics analysis was performed on BM and PB cell pellets, yielding 73 and 91 lipids, respectively, across fifteen different lipid subclasses (Figure 1E-F). Significant variance in overall lipid abundance was evident at acute primary infection, although both compartments displayed similar patterns of upregulation on the level of individual lipids (Figure 1G-H). Phosphatidylcholines and phosphatidylinositols were the major lipid types upregulated in both compartments. These results are also consistent with the metabolomics measurements. Pathway analysis using mummichog (Li et al., 2013) and MetaCore indicated prominent changes in metabolism of bile acids and immune regulators (e.g., prostaglandins, leukotrienes, and lipoxins) during acute infection (**Table S3**). This is consistent with results from multiple other data types, including changes in transcripts for cytosolic phospholipase A2 (cPLA2, which catalyzes production of arachidonic acid) and cyclooxygenase 1 (COX-1, which converts arachidonic acid to prostaglandins), both of which were significantly upregulated in PB during acute infection (Figure S1D-E), as well as the measured increase in lipid precursors of arachidonic acid described above.

### Integrative Multi-omic Analysis Provides Improved Identification of Differences between Infection Time points in PB and BM

Based on the improvement in PCA clustering when all data types were naively integrated, we sought another method to directly integrate the various data types to exploit their potential overlaps in molecular patterns. We used a machine learning approach based on the identification of mutual information between data types to accomplish multi-omic integration. Mutual information has been widely used as a similarity metric when inferring biological networks (Aguilar et al., 2012; Kim et al., 2010; Margolin et al., 2006), enabling better identification of nonlinear relationships than many correlation-based approaches. This advantage is particularly important given the expected nonlinear and complex relationships across different omics data types, which may underlie critical regulatory mechanisms (Becavin and Benecke, 2011). Here, we used a well-established mutual information-based algorithm, Context Likelihood of Relatedness (CLR) (Faith et al., 2007), to develop networks consisting only of features with high mutual information across multiple data types, thus capturing events whose impacts span molecular layers.

CLR was performed pairwise on the different data types (BM or PB transcriptomics, BM or PB lipidomics, and metabolomics), and the resulting compartment-specific networks were then used as the basis for further analyses. Features having no significant mutual information with variables in any other data type were removed from the network and the subsequent analyses. PCA of individual data types using the reduced variable sets (Figure 3C) showed significantly improved separation between the sample classes in almost all cases. Moreover, PCA on variables across all data types in this multi-omics network provided remarkable separation between pre-infection, acute infection, and relapse time points in both BM and PB (Figure 3D). This separation is particularly noteworthy because it is essentially unsupervised: selection of variables is being performed based on a completely separate criterion (mutual information across data types). This suggests that variables with significant relationships to variables in other data types are excellent indicators of, and may even be causes of, observable phenotypes.

### Compartment-specific Multi-omics Relatedness Networks Capture the Systems-Level Landscape and Dynamics of Host Responses to Malaria

For further biological interpretation and characterization of the multi-omics network we derived, we used the transcriptional data as the basis for segmentation of the network and annotation of the resultant modules. Specifically, we identified all transcriptional pathways that were enriched in the overall derived multi-omics network, and used those to define smaller Multi-Omics Relatedness Networks (MORNs). These MORNs consisted of the network’s genes from a given enriched pathway, together with all of the lipids and metabolites with which they had significant mutual information. We constructed atlases of these MORNs for both BM (Figure 4A) and PB (Figure S2). While most enriched transcriptional pathways were compartment-specific, both compartments showed many enriched immune and cytokine signaling pathways (Figure 4B). Interferon-gamma (IFN-γ) and tumor necrosis factor (TNF) are well-known factors that contribute to the pathogenesis of infection (King and Lamb, 2015; Meding et al., 1990; Villegas-Mendez et al., 2012), and both genes were upregulated in acute, but not relapsing, infection time points. Concomitantly cytokine levels of IFN-γ and TNF were elevated at the acute point of infection returning close to baseline after the peak (Figure 4C-D). These responses were accompanied by the induction of interleukin (IL)-12 (Figure 4E-F), a key cytokine driving IFN-γ responses, as well as strong upregulation of IFN-γ-induced protein (IP)-10 (Figures 4G and 4H), a key chemokine induced by IFN-γ (Freitas do Rosario et al., 2012) also known as CXCL10. Altogether this demonstrates that, as expected, in these experiments *P. cynomolgi* induced a strong type 1 inflammatory response associated with IFN-γ. These observations are consistent with human studies (Jagannathan et al., 2015; Walther et al., 2009). As such, the transcriptome associated with this inflammatory response can be investigated to discover the importance of other immunological processes that may be occurring in *Plasmodium* infections.

**Figure 4.**
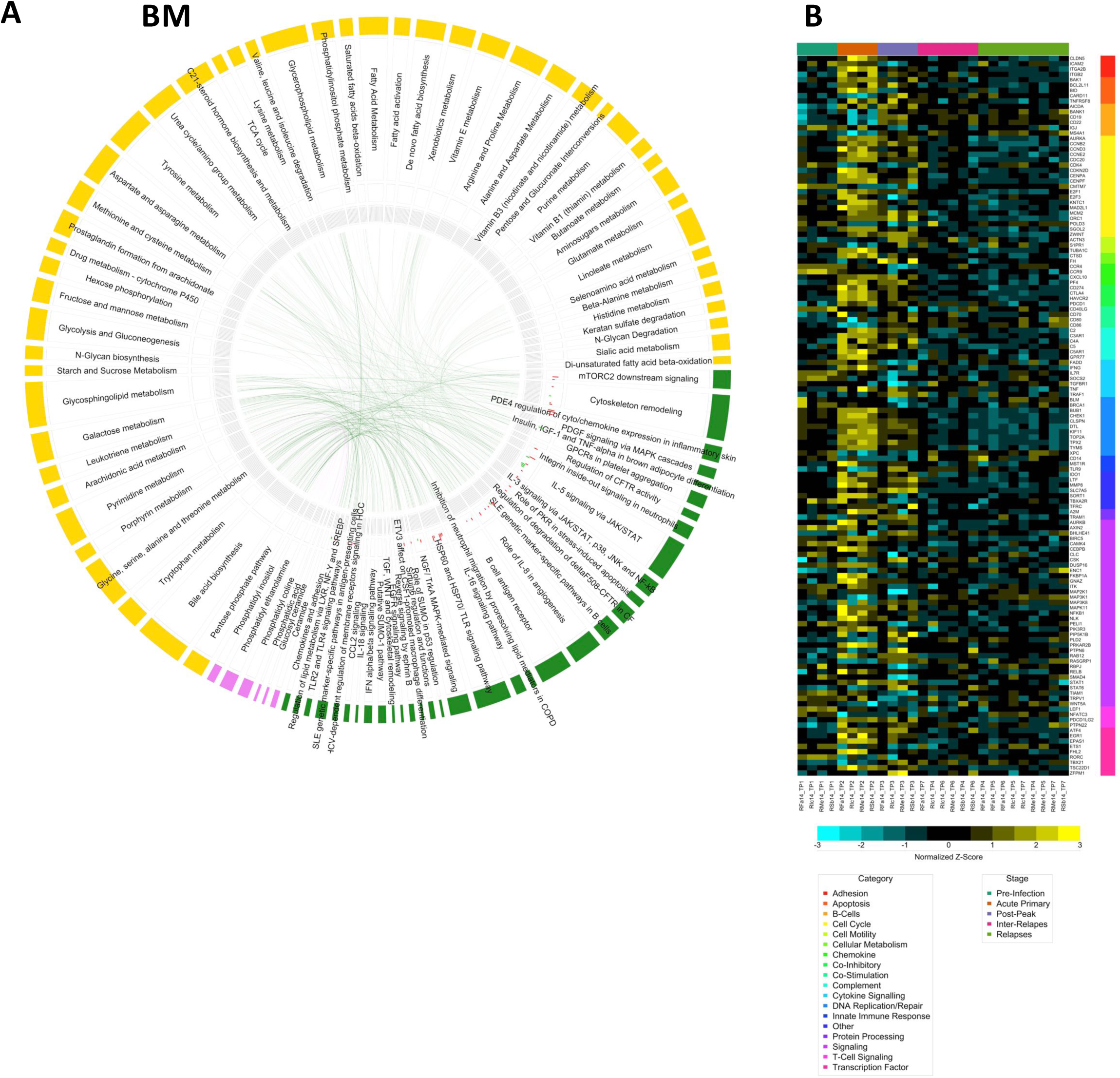
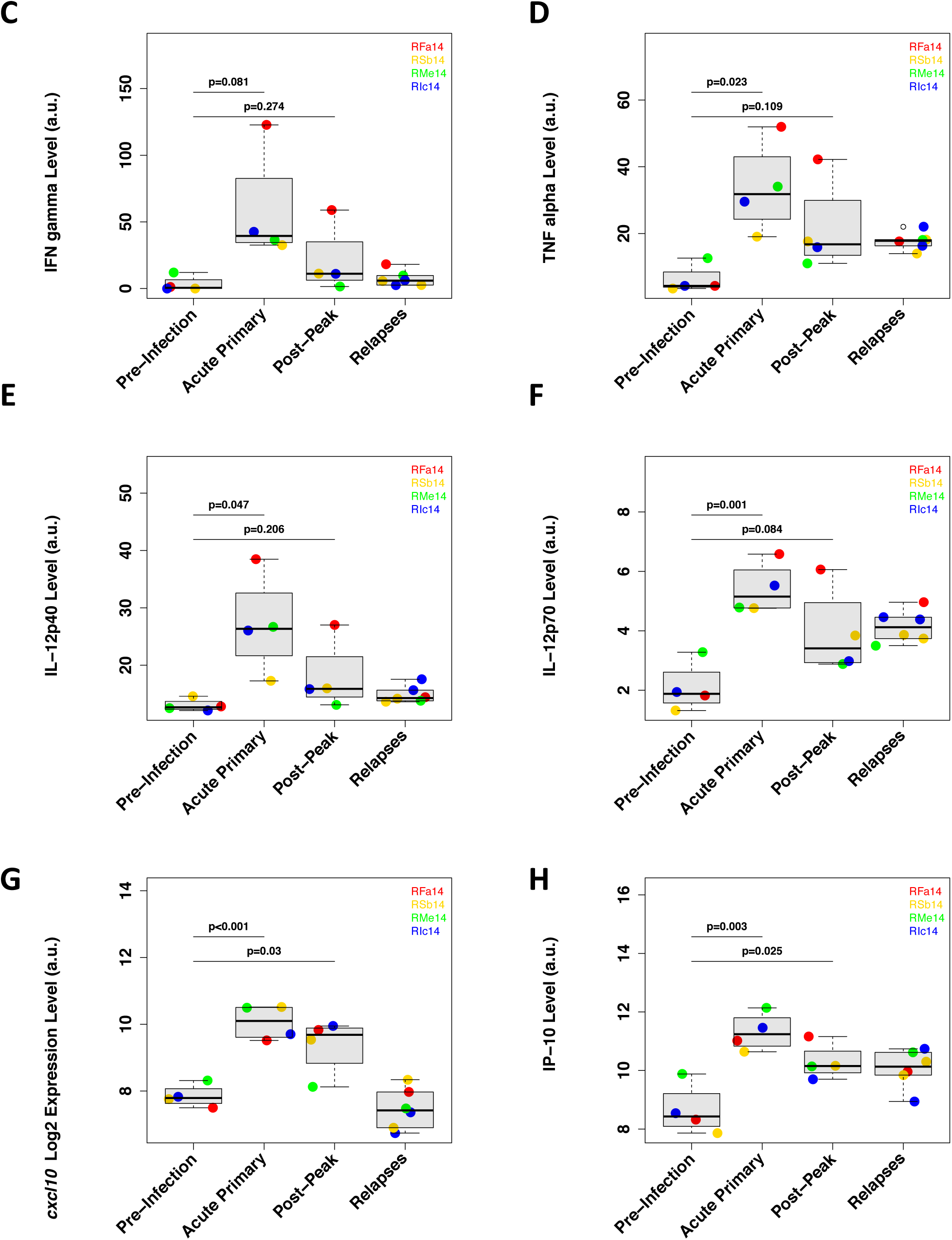
Atlas of Multi-Omics Relatedness Networks (MORNs). (A) Atlas of MORNs identified in BM. Segments in the outer ring of the circle represent metabolic pathways (yellow), gene pathways (green) and types of lipids (pink), with size corresponding to numbers of nodes or molecules in the pathway/category. Edges of the network (connecting lines) are defined by having significant mutual information scores between connected nodes. (B) Enriched immune and cytokine signaling pathways found in both PB and BM. (C) Upregulation of Interferon-gamma (IFN-γ) plasma levels at acute infection. (D) Upregulation of tumor necrosis factor (TNF) plasma levels at acute infection. (E-F) Upregulation of IL-12 plasma levels at acute infection. (G-H) Upregulation of IP-10 transcriptional (G) and plasma (H) levels at acute infection.

Of the top DEGs in our MORNs, several pathways previously shown to be important in regulating immune function were enriched in both compartments, including the mechanistic target of rapamycin (mTOR) signaling axis and the sirt6 pathway (Bauer et al., 2012; Weichhart et al., 2015). MORNs related to immune cell function were also positively correlated with cytoskeletal remodeling and cell cycle regulation, possibly as a result of the extensive expansion and migration that occurs in the immune cells participating in the immune response. We observed noticeable correlation structure across BM and PB MORNs (Figure S3A-B). The PB MORNs formed two large clusters that were negatively correlated with each other, with extensive enrichment for immune response functions (immune cell signaling pathways and cytokine signaling pathways).

Changes in correlation structure of variables within each MORN as a function of time (Figures S4A and S4B) hint at the potential for regulatory networks to be reshaped during acute infection in a way that exerts long-term effects at the time of the relapse. The trajectories of the MORNs (expressed as their eigenfeature across pre-infection, acute infection, post-peak, and relapse time-points) can often be functionally validated via other measurement modalities (Figure S4C-D). However, given the complexities that arise from capturing only the peripheral component of the immune response, these do not always correlate. In addition to immune transcriptomic profiles, there was a significant overlap of enriched metabolic pathways in BM and PB MORN atlases, potentially reflecting systemic involvement of metabolic regulation in malaria.

### Relating MORNs to Host Phenotypes

To identify MORNs of interest for more detailed analysis and validation, we sought to identify correlations between MORNs (via their eigenfeatures) and phenotypic measurements (immunophenotyping, plasma cytokine profiles, and clinical traits). The eigenfeature of each MORN was used to assess correlations with these phenomics measurements, using pairwise biweight midcorrelation. Figure 5 contains a representative subset, and Figure S5 shows all significant interactions with clinical traits. Full correlation networks are shown in Figure S6.

**Figure 5.**
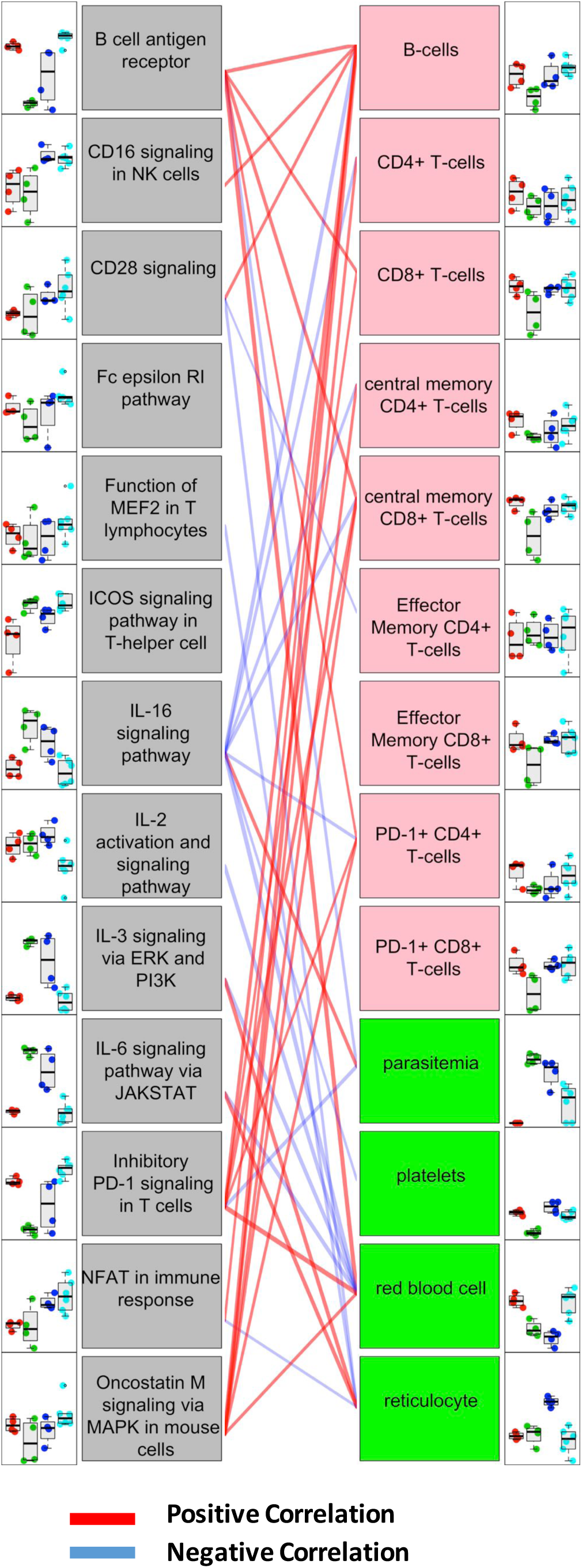
A representative subset of identified correlations between MORNs and phenotypic features. Trajectories of the eigenfeatures for MORNs are connected to phenotypic feature profiles with which they are positively (red) or negatively (blue) correlated.

Among PB MORNs, some of the most significant positive correlations with clinical traits were between parasitemia (which peaked at acute infection and had much lower, but non-zero, levels during relapse) and MORNs involving cytokine signaling (Figure S5A). Some MORNs were suggestive of function (e.g., MORNs related to immune cell functions such as PD-1 signaling in T cells that were negatively correlated with parasitemia). However, given the dual nature of some of these molecules and the peripheral location of PBMCs that may not necessarily reflect organ-specific immune responses, such correlations with the phenotype of infection remains to be tested.

### Applicability of NHP Malaria MORNs to Human Vivax Malaria

To assess the applicability of MORNs (and insights derived from them) from the NHP model of *P. cynomolgi* infection to human vivax malaria, we tested whether changes in host transcriptional pathways associated with the MORNs identified here could be recapitulated using a recently published human PB transcriptomics dataset from *P. vivax* patients (GSE67184) (Rojas-Pena et al., 2015). 735 of 6,154 genes were differentially expressed at the time of diagnosis, yielding 262 significantly enriched curated pathways (FDR < 0.05). Out of 47 transcriptional pathways identified by PB MORNs from the NHP model, enrichment was found for 25 of those pathways in the human transcriptomics dataset, including the PD-1 signaling pathways and several malaria-relevant interleukin signaling pathways (**Table S4**). The meta-enrichment in the human transcriptomic dataset of pathways associated with NHP MORNs was significant via a hypergeometric test (p < 0.01), supporting the utility and translational potential of the MORN approach and the NHP model system to study malaria. In fact, a high degree of concordance between NHP and human fold-change levels was found for the transcriptional gene sets represented by MORNs. All three of the gene sets with at least 15 genes measured in both the NHP and human datasets (including PD-1 signaling) showed highly significant correlations (Figure 6A), while other gene sets with less overlap between the datasets showed similar trends toward concordance. Thus the transcriptome of this human vivax data set support the utility of the *P. cynomolgi* model in rhesus macaques for understanding host-pathogen interactions in *P. vivax* infection.

**Figure 6.**
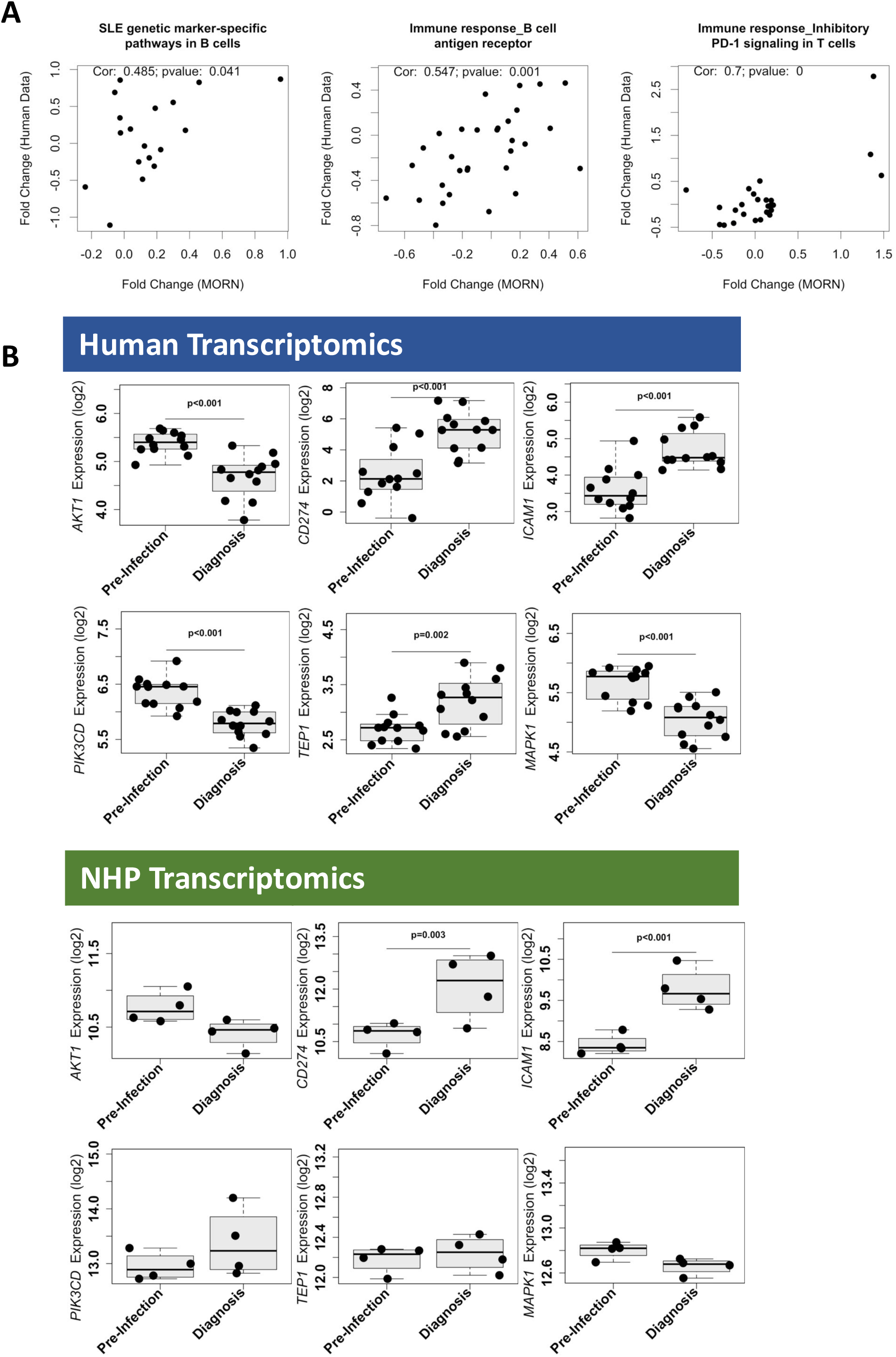
Assessment of NHP malaria MORNs in human vivax malaria. (A) Gene sets with at least 15 genes measured in both the NHP and human datasets showed high, significant correlations for expression levels in these two datasets. The x-axis represents the log_10_ fold change of gene expression in NHP model, and the y-axis represents the log_10_ fold change of gene expression in human data. (B) Transcriptional profile comparisons of PD-1 signaling pathway genes between NHP model and human data.

The PD-1 pathway had the most significant correlation between the NHP and human transcriptional measurements. Pathway analysis on DEGs identified in the *P. vivax* patient transcriptomics data showed enrichment of PD-1 signaling at FDR < 0.05. Most of the enriched transcripts from the PD-1 signaling pathway showed similar expression trends as in the *P. cynomolgi* – rhesus macaque model (Figure 6B). Integration of multi-omics data thus helped reveal the potential generalizability of PD-1 involvement in *Plasmodium* infection, highlighting the power of using a multi-omics approach in the identification of pathways of importance in infection models.

## Discussion

This unprecedented set of longitudinal NHP multi-omics measurements all from the same samples, coupled with a novel computational method to directly integrate these datasets, constitute a unique resource for research on malaria caused by *P. cynomolgi* and its sister species, *P. vivax*. Using this resource, we constructed the first global atlas of multi-omics subnetworks in PB and BM, two important immune compartments reflecting outcomes and pathogenesis of *Plasmodium* infection. This approach revealed molecular signatures for primary *Plasmodium* infection in rhesus macaques, identified measurements capable of distinguishing acute infection and relapse, and identified pathway involvement that is also recapitulated in prior studies of human malaria.

Differential analysis of individual transcriptomics, metabolomics, and lipidomics datasets showed significant alterations of transcriptional and metabolic programs related to inflammation and immune regulation during acute infections that are consistent with a number of reported observations (Auclair et al., 2014; Haque et al., 2001; Sarangi et al., 2014; Sinha et al., 2010; Zago et al., 2012). Altered pathways were related to two axes of immune regulation: those modulating cell proliferation through the cell cycle and those controlling immune function (including cellular immunity). The extensive enrichment of cytokine signaling pathways observed in both compartments was corroborated by increased levels of cytokines measured in the plasma, while lipid, transcriptional, and metabolic data together supported the importance of pro-inflammatory lipid species during acute infection.

Interrelationships among omics layers can provide unique insights about processes and networks that span multiple molecular layers and impact immune system regulation. To address this question, we used a mutual information-based network analysis approach to directly integrate multi-omics data types, rather than post hoc identification of ontological and interpretation similarities between individual omics analyses. This approach enabled improved ability to distinguish between infection time points. The selection of variables for these analyses was based solely on mutual information across molecular layers without information about time points, underscoring the robustness of this approach. Compartment-specific atlases of multi-omics networks, charting some of the complexity of the multi-level landscape underlying malarial pathogenesis and immunity to parasite infection, were also generated to direct future studies in this area.

Importantly, the analysis revealed a strong Th1-biased inflammatory response to *P. cynomolgi* infection as expected from previous studies of mouse and human *Plasmodium* infections. Two signaling pathways known to be involved in immune memory, mTOR and PD-1 signaling, were enriched in the PB MORNs and were negatively correlated with parasitemia. However, above the identification of novel molecules potentially involved in malaria that can now be taken for further study, our approach to multi-omics integration also identified a subset of molecules and pathways that are unique to relapse. Overall immune, transcriptional and other responses during relapse were quite muted, highlighting the importance of finding unique relapse responses in an unsupervised fashion. Although the transcriptomic signatures at relapse may, in part, be muted due to the presence of hemoglobin in the samples, the transcriptome of hemoglobin-depleted samples from this study has also been carried out and is included in the data repository for further comparison.

Relapse is a major bottleneck for malaria treatment and elimination of *P. vivax* due to the dormant liver form of the parasite. The *P. cynomolgi* NHP model provides a platform to study relapse (Joyner et al., 2015; Joyner et al., 2016). Our analysis of longitudinal infections provides insight on key signaling pathways that may form the basis for novel strategies and drug targets for relapse treatment and prevention via modulation of host immune memory. We expect the datasets presented here and the multi-omic integration strategy we have used to be a valuable resource for the malaria and non-human primate research communities, helping enable investigations of the pathogenesis of malaria and the complexity of the host response to this parasite.

---

Figure S1. Global characterization of omics profiles. (A) Acute infection was characterized by the broadest scope and greatest magnitude of changes in omic profiles. Features were clustered across all omics data. Samples were organized by infection time points (Pre-Infection, Acute, Post-Peak, Inter-Relapses, Relapses). Heatmap includes transcriptomics, metabolomics, and lipidomics from PB and BM. Standardized data are plotted. Color bars on left of the figure represent types of omics features: transcripts from PB (HostPB, red), transcripts from BM (HostBM, pink), lipids from PB (LipidPB, blue), lipids from BM (LipidBM, cyan), metabolites from serum (Metabolomics, gold).

(B) Differential expression of genes at relapse in PB.

(C) Differential expression of genes at relapse in BM.

(D)Boxplots of gene expression of pla2g4c at acute primary infection.

(E) Boxplots of gene expression of ptgs1 at acute primary infection.

Figure S2. Atlas of Multi-Omics Relatedness Networks (MORNs) in PB. Segments in the outer ring of the circle represent metabolic pathways (yellow), gene pathways (green) and types of lipids (pink), with size corresponding to numbers of nodes or molecules in the pathway/category. Edges of the network (connecting lines) are defined by having significant mutual information scores between connected nodes.

Figure S3. Clusters of MORNs. (A) Clustering of MORNs identified in PB based on pairwise correlations. Red indicates positive correlation, blue indicates negative correlation.

(B) Clustering of MORNs identified in BM based on pairwise correlations. Red indicates positive correlation, blue indicates negative correlation.

Figure S4. Dynamics of MORNs. (A) Dynamics of MORNs in PB. Each contour plot represents the distribution of loadings of each variable in the MORN across the first two principle components of the data for those variables at a given infection point and across all replicates. Changes in contour plots indicate changes in relative importance of MORN variables across different infection points.

(B) Dynamics of MORNs in BM. (C-D) Heatmaps of eigenfeatures of PB (C) and BM (D) across infection points.

Figure S5. Relationships between MORNs with phenotypic data. (A) Correlation between MORNs (PB) and clinical traits. The colors of the boxes are scaled with the value of correlation coefficients using bicor analysis, ranging from green (r = −1) to red (r = 1). Correlations with significant pvalue<0.05 are shown with numbers indicating correlation coefficient (first line) and pvalue (second line).

(B) Correlation between MORNs (BM) and clinical traits.

(C) Correlation between MORNs (PB) and immune profiles.

Figure S6. Centrality Atlas constructed in BM (A) and in PB (B). Network icons represent MORNs; gold spheres represent adaptive immunological profiles; green spheres represent innate immunological profiles; purple rectangles represent cytokine profiles; pink circle represent clinical traits. Solid lines between nodes represent significant positive correlation; dotted lines represent significant negative correlation.

