## Supplementary figures and images for "Multi-omics Integrative Analysis of Acute and Relapsing Malaria in a Non-Human Primate Model of *P. vivax* infection"

### Supplementary Figures

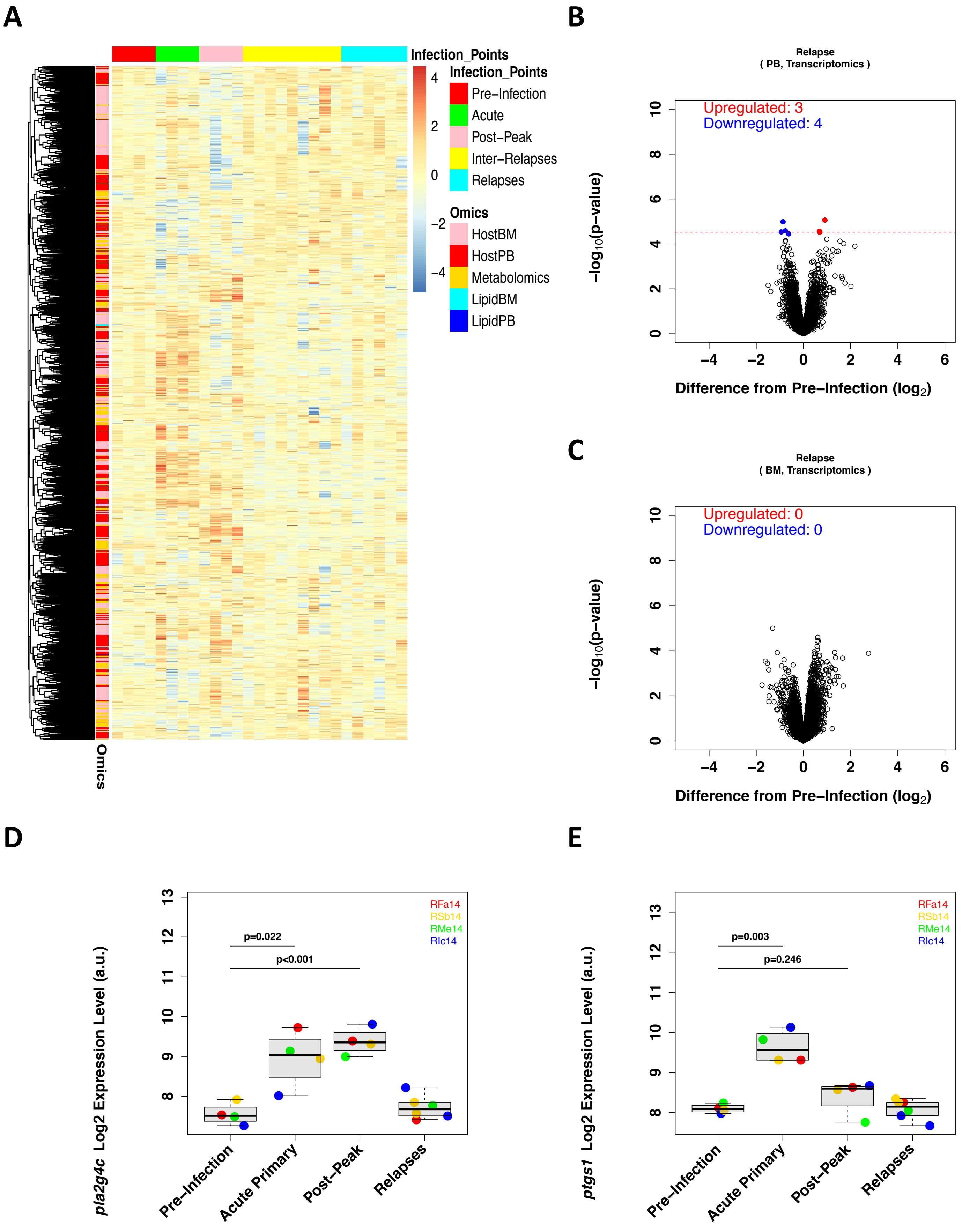

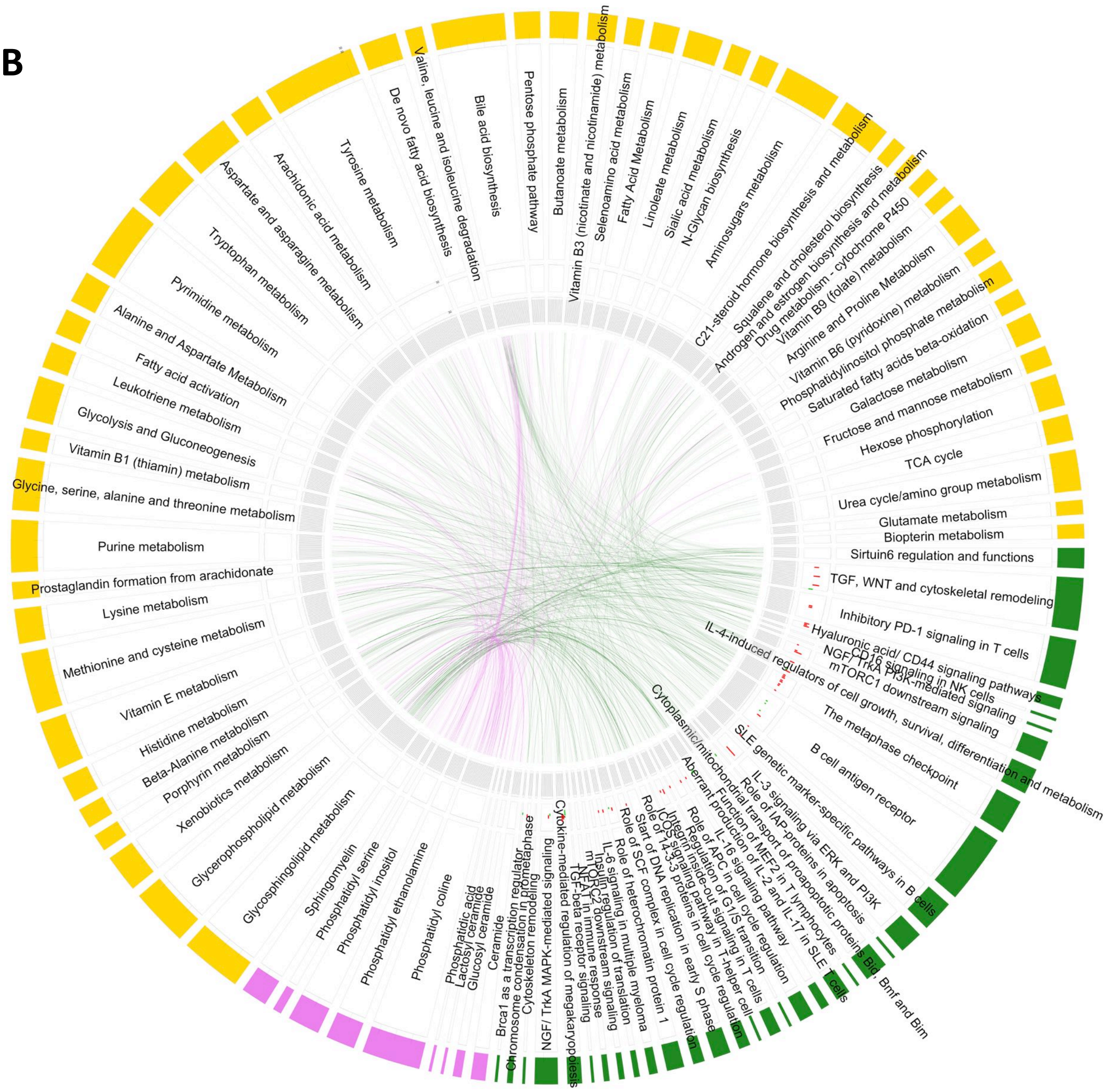

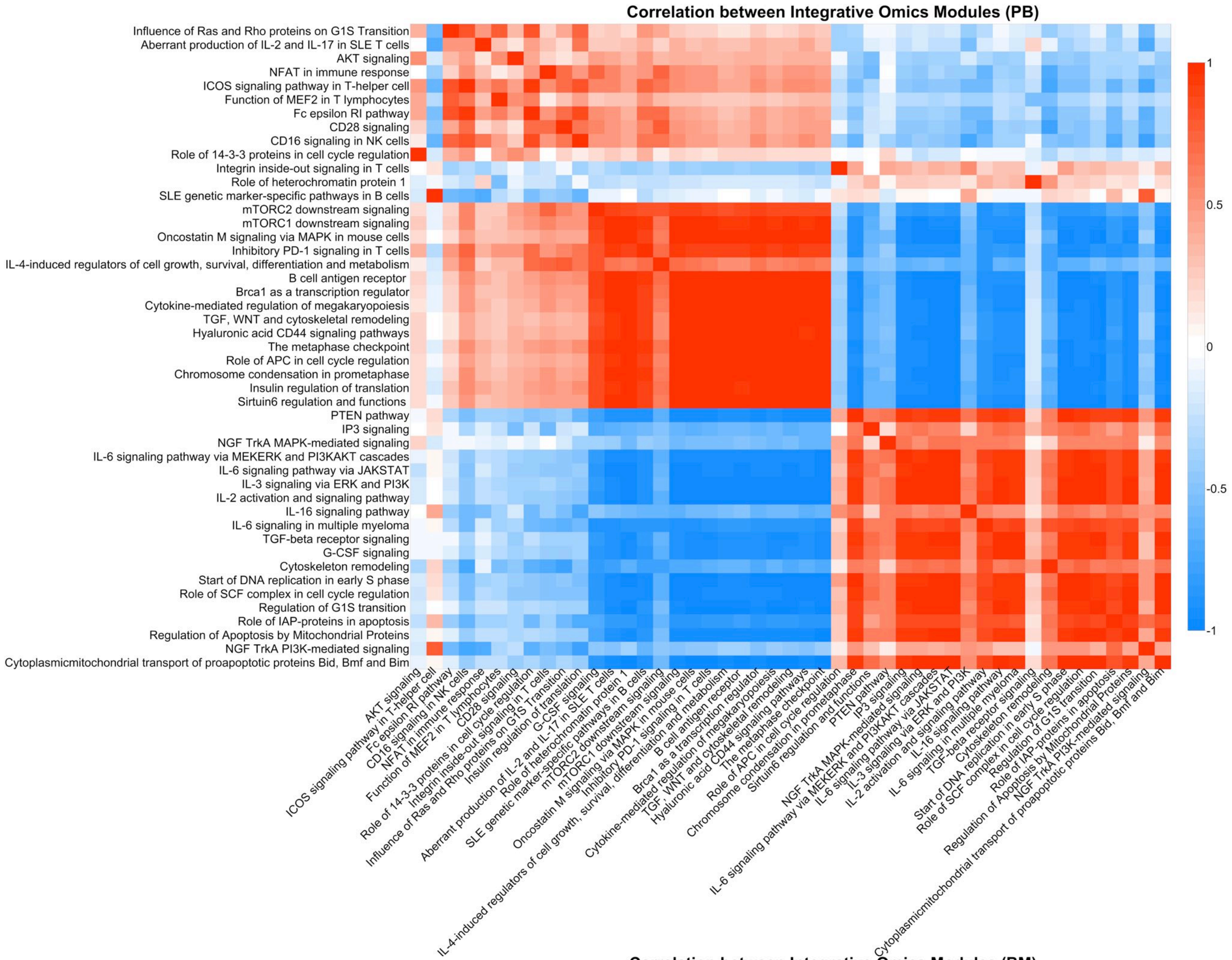

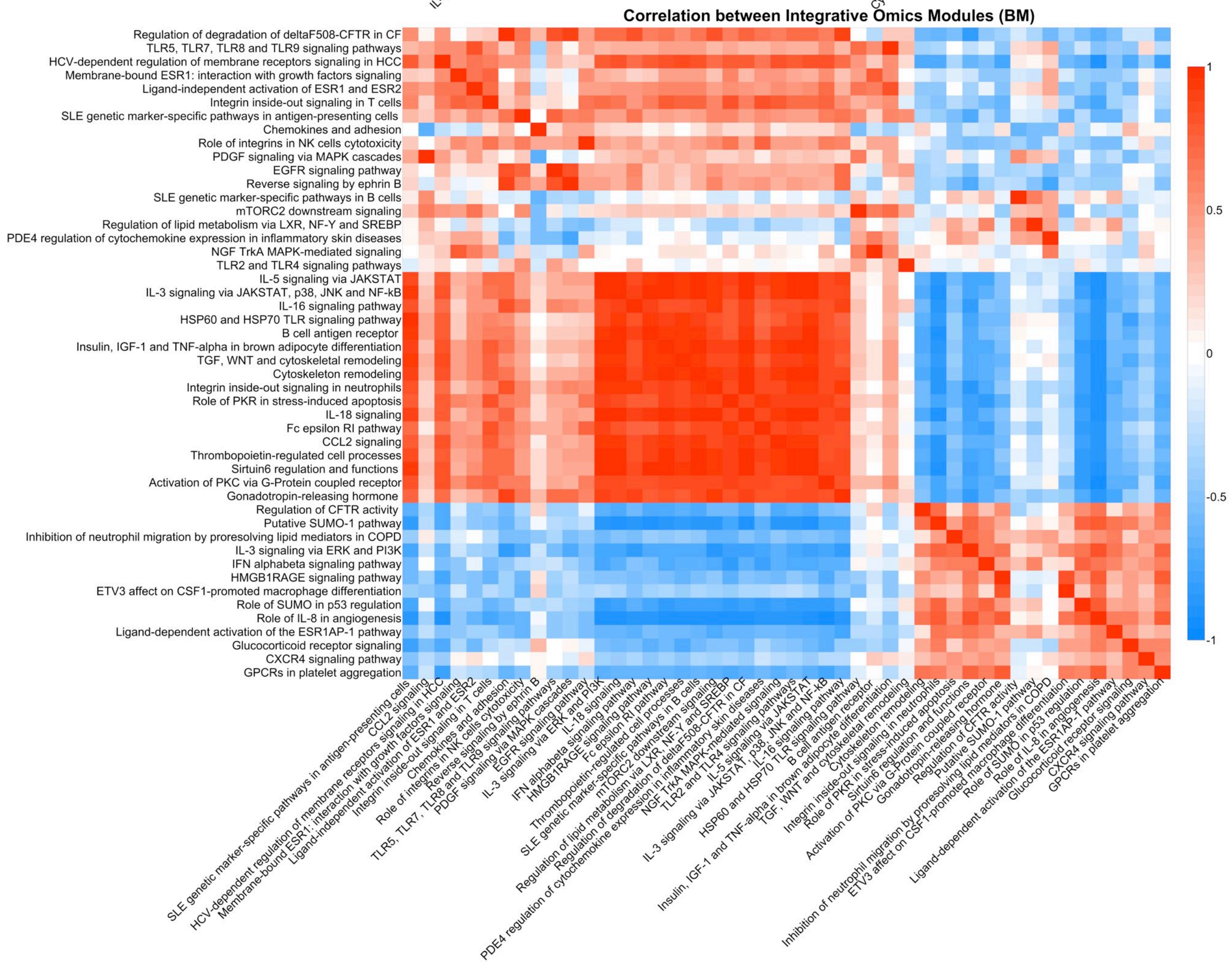

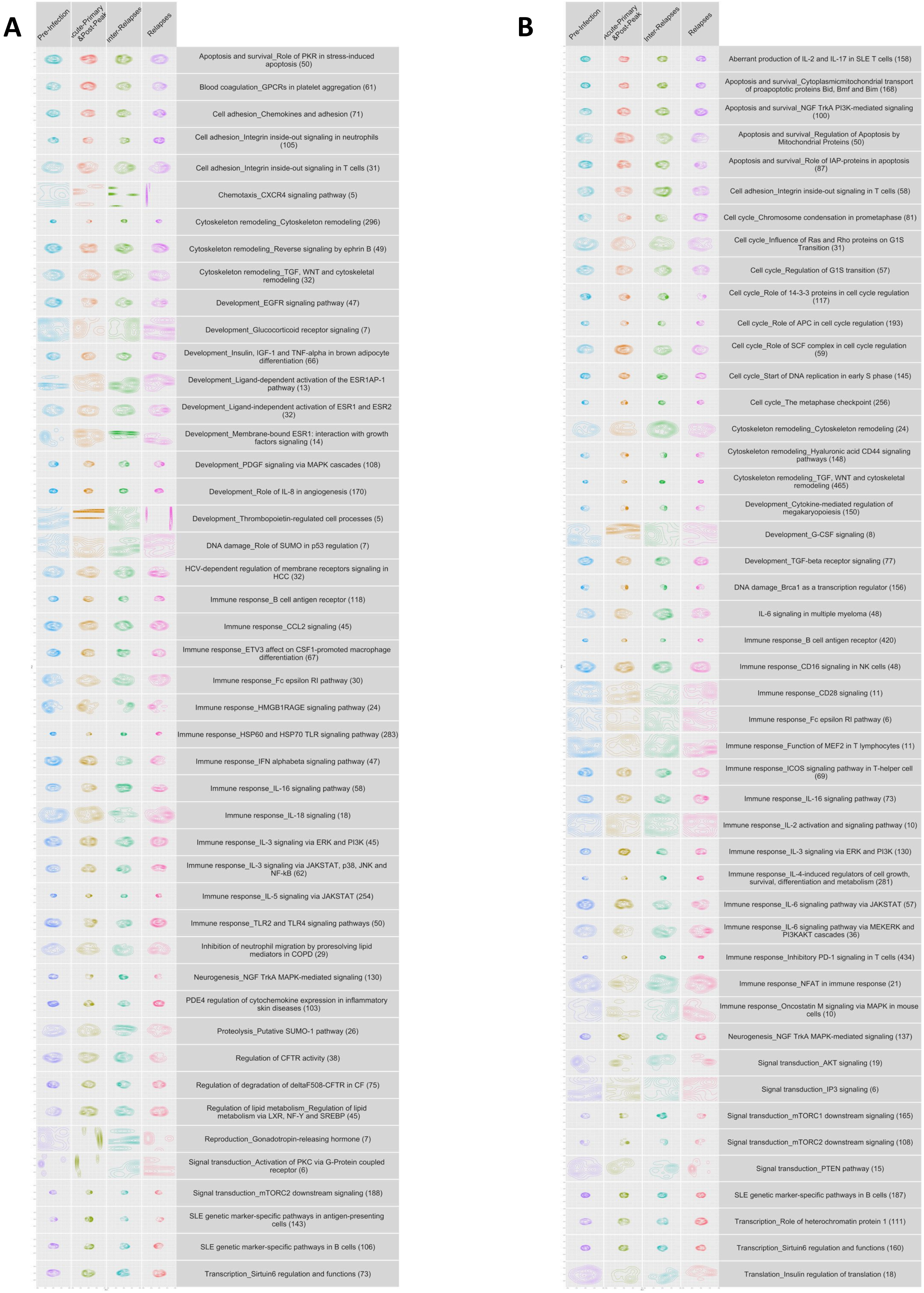

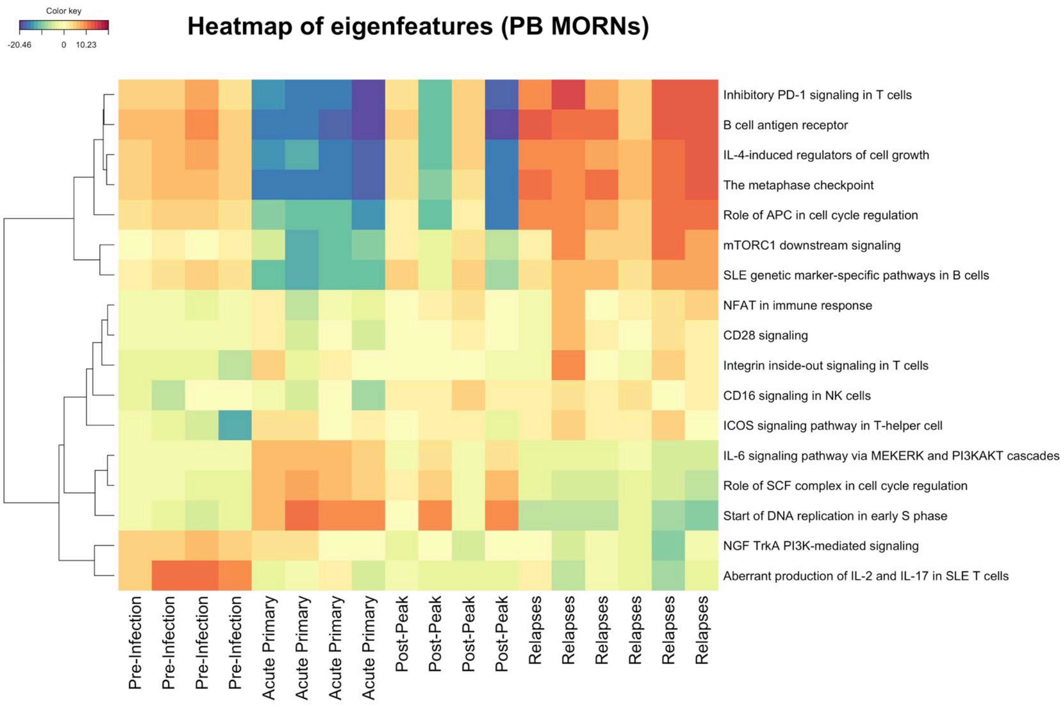

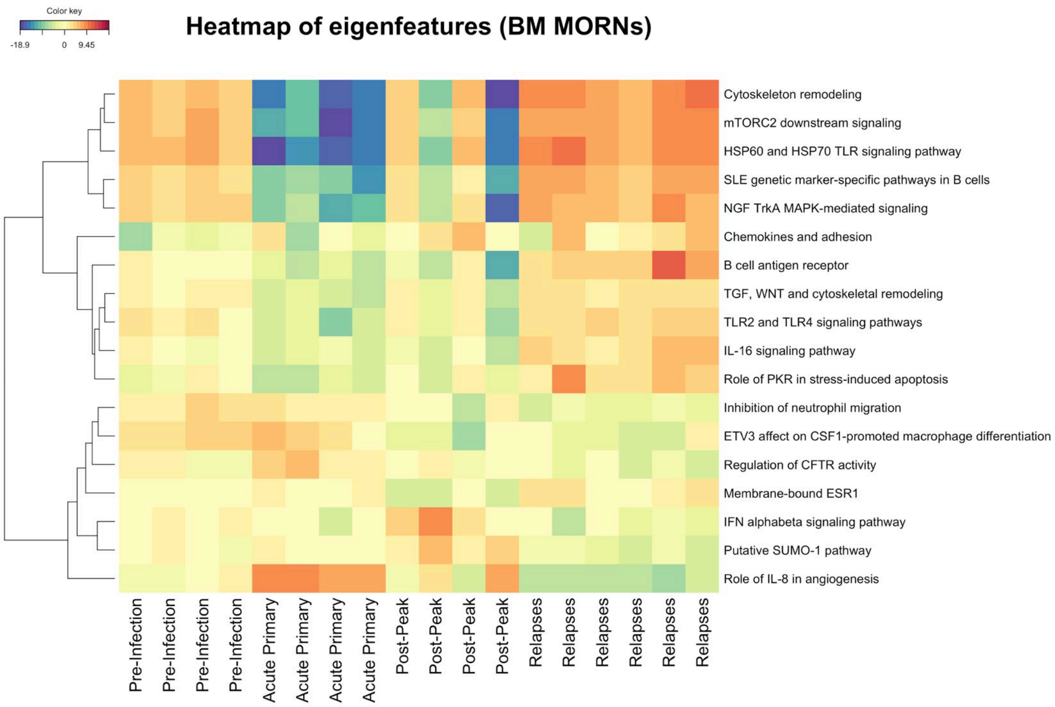

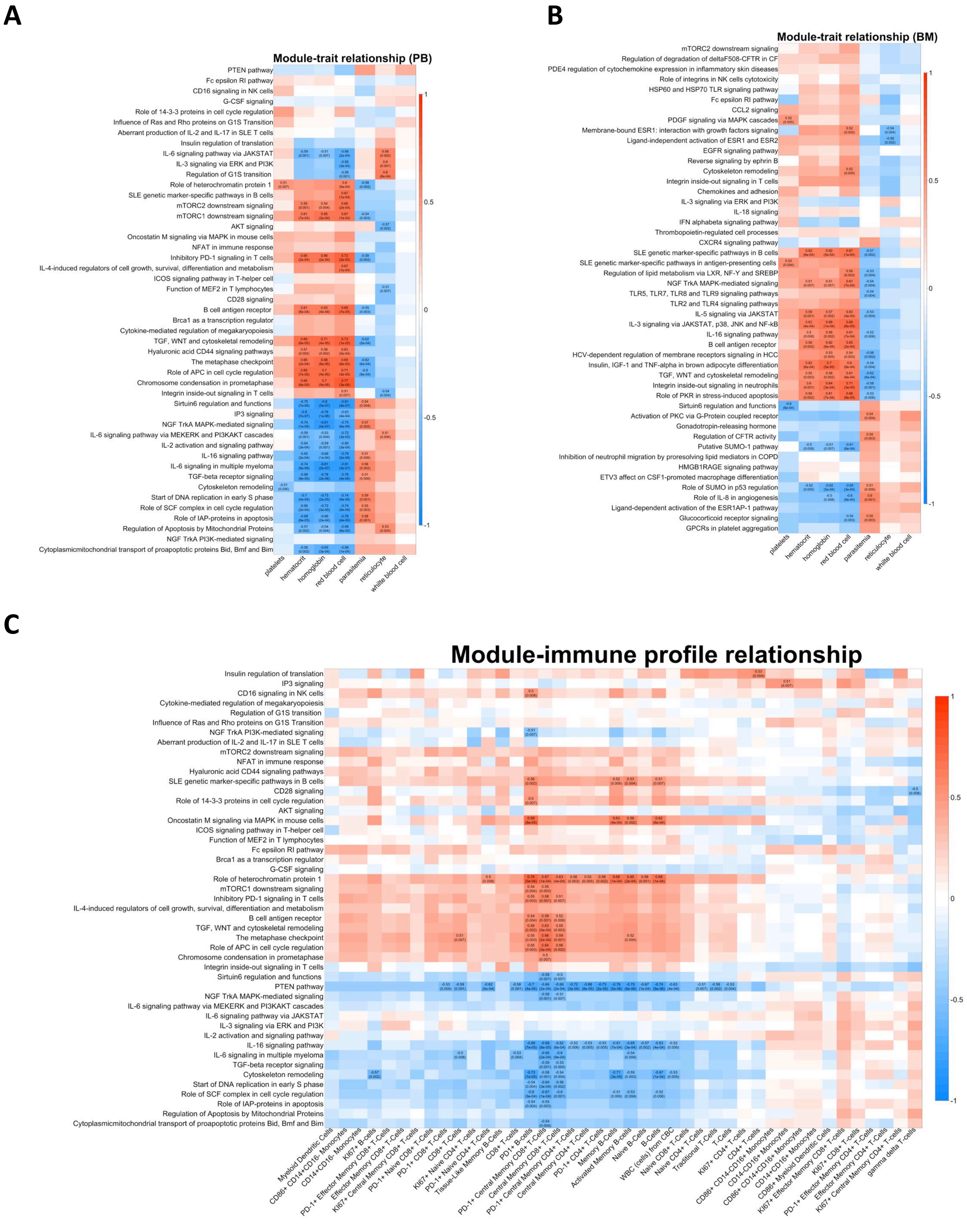

A

BM

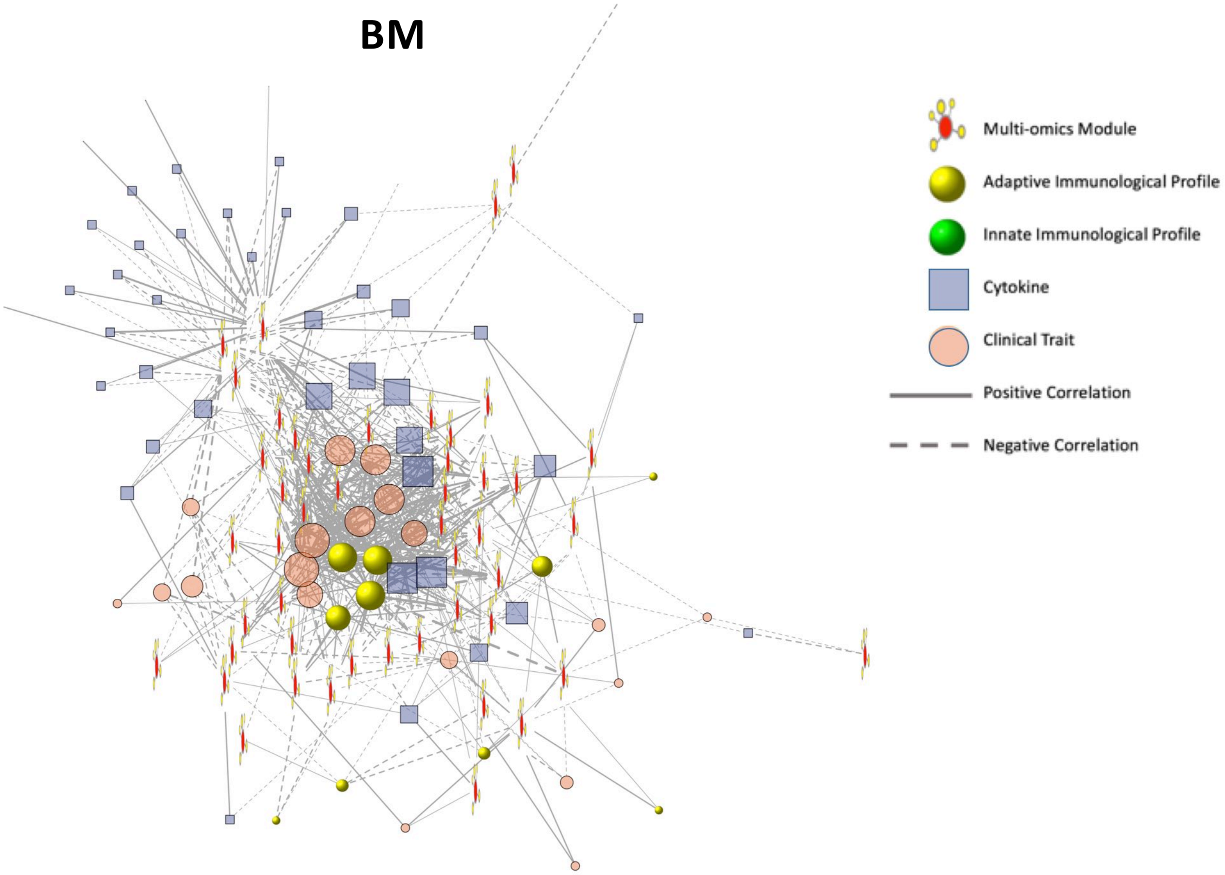


B

PB

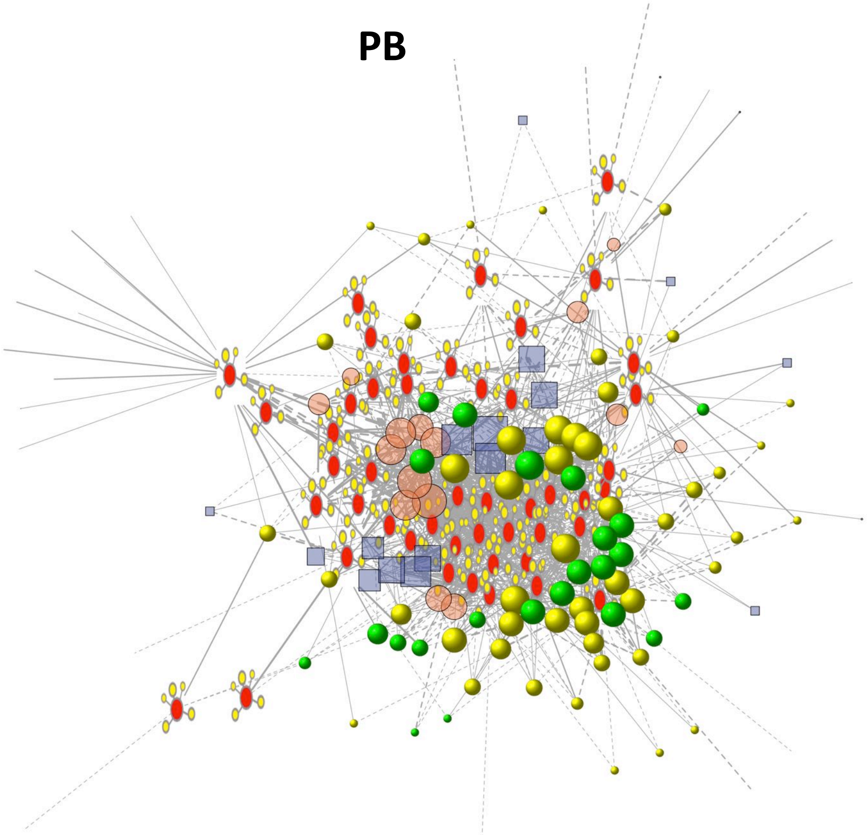

**Figure S6**
